# G_i/o_ protein–coupled receptor inhibition of beta-cell electrical excitability and insulin secretion depends on Na^+^/K^+^ ATPase activation

**DOI:** 10.1101/2022.02.10.479802

**Authors:** Matthew T. Dickerson, Prasanna K. Dadi, Karolina E. Zaborska, Arya Y. Nakhe, Charles M. Schaub, Jordyn R. Dobson, Nicole M. Wright, Joshua C. Lynch, Claire F. Scott, David A. Jacobson

## Abstract

G_i/o_ protein-coupled receptors (G_i/o_-GPCRs) limit pancreatic islet insulin secretion by decreasing β-cell Ca^2+^ entry, which is essential for maintenance of glucose homeostasis. However, the G_i/o_-GPCR signaling mechanism that mediates inhibition of human islet hormone secretion has not been identified. Here we demonstrate that G_i/o_-GPCRs cause hyperpolarization of the β-cell membrane potential through activation of Na^+^/K^+^ ATPases (NKAs) in mouse and human islets. Stimulation of G_i/o_-coupled somatostatin or α2-adrenergic receptors induced oscillations in β-cell NKA activity, which resulted in islet Ca^2+^ fluctuations. Selective induction of β-cell G_i/o_ signaling with a chemogenetic G_i/o_-GPCR also activated NKAs and initiated islet Ca^2+^ oscillations, suggesting that β-cell G_i/o_-GPCRs tune pulsatile insulin secretion. Furthermore, intra-islet paracrine activation of β-cell G_i/o_-GPCR signaling and NKAs by δ-cell somatostatin secretion slowed Ca^2+^ oscillations, which decreased insulin secretion. G_i/o_-GPCR-mediated oscillations in β-cell membrane potential and Ca^2+^ were dependent on NKA phosphorylation by Src tyrosine kinases; an effect that was mimicked by stimulating islet insulin receptor tyrosine kinases. Whereas β-cell NKA function was completely inhibited by cAMP-dependent PKA activation. Taken together, these data reveal that NKA-mediated hyperpolarization of β-cell membrane potential serves as the primary and conserved mechanism for G_i/o_-GPCR control of electrical excitability, Ca^2+^ handling, and insulin secretion.

## INTRODUCTION

Pancreatic β-cell glucose-stimulated insulin secretion (GSIS) is essential for maintenance of euglycemia (1, 2), and as Ca^2+^ entry is required for GSIS, mechanisms that control β-cell Ca^2+^ handling are critical regulators of blood glucose homeostasis (3–7). It was discovered more than 50 years ago that G_i/o_-coupled G protein-coupled receptors (GPCRs) play a critical role in limiting insulin secretion in-part by decreasing β-cell electrical excitability and subsequent Ca^2+^ influx (8–10). However, the exact mechanism(s) of G_i/o_-coupled GPCR control of β-cell electrical excitability, Ca^2+^ handling, and insulin secretion remain poorly understood.

β-cells express numerous G_i/o_-coupled GPCRs such as somatostatin receptors (SSTRs), α2A-adrenergic receptors (ADRs), and D2-like dopamine receptors (DRDs) (11–16). Treatment with G_i/o_-coupled GPCR ligands (i.e. somatostatin (SST), adrenaline, or dopamine) activates hyperpolarizing currents and reduces intracellular cAMP ([cAMP]_i_) levels, which results in decreased intracellular Ca^2+^ ([Ca^2+^]_i_) and diminished insulin secretion (16–21). As insulin secretion is inhibited by G_i/o_-coupled GPCRs, these signals are critical for preventing excessive insulin secretion under hypoglycemic as well as stimulatory conditions (22). Indeed, loss of intact α2-ADR signaling leads to a drop in blood glucose levels under fasting and fed conditions due to elevated insulin secretion (23), while ADR agonists attenuate GSIS (24, 25). Intra-islet communication is also mediated via G_i/o_ signaling (i.e. SST is secreted by δ-cells while dopamine is secreted by β- and α-cells) (26–30), which tunes β-cell Ca^2+^ handling and insulin secretion. Thus, inhibition of islet G_i/o_-coupled GPCRs with pertussis toxin, also known as islet activating protein (31), significantly stimulates hormone secretion, highlighting the importance of G_i/o_-coupled GPCRs in regulating islet function. As numerous G_i/o_-coupled GPCRs control physiological β-cell function, perturbations in these pathways impair β-cell GSIS and are in some instances associated with increased risk of developing diabetes (14, 32–34). For example, glucose-stimulated SST secretion is blunted during the pathogenesis of type 2 diabetes (T2D), which diminishes SSTR-mediated control of β-cell function (33). Moreover, polymorphisms that increase α2A-ADR expression result in increased risk of developing T2D due to suppression of GSIS (14, 32, 34). Taken together, these findings strongly suggest that G_i/o_ signaling plays a key role in regulating β-cell Ca^2+^ handling and insulin secretion; however, the underlying mechanism has not been conclusively identified for more than half a century.

β-cell *V*_m_ hyperpolarization is predominantly mediated by K^+^ efflux; thus, it has been generally accepted that G_i/o_-coupled GPCR signaling activates an outward K^+^ conductance. However, ATP-sensitive K^+^ (K_ATP_) channels can be ruled out as the source, because activation of G_i/o_-coupled GPCRs induces β-cell *V*_m_ hyperpolarization in the presence of sulfonylureas and in K_ATP_ channel-deficient islets (21, 35). As is the case in numerous other tissues, G_i/o_ signaling-induced β-cell *V*_m_ hyperpolarization has been widely ascribed to activation of G protein-gated inwardly-rectifying K^+^ (GIRK) channels (16, 36). RNA sequencing studies show that both mouse and human β-cells express low levels of GIRK channel transcripts (predominantly *KCNJ6,* the gene encoding GIRK2) (11, 12, 16), and immunofluorescent staining of mouse pancreatic sections confirms expression of GIRK channel proteins in β-cells (36). While G_i/o_ signaling activates robust inwardly-rectifying K^+^ currents in other cell types where GIRK channels are expressed (37), currents with GIRK-like characteristics have not been reproducibly observed in primary β-cells. Moreover, there are conflicting reports detailing the effect of GIRK channel inhibitors on β-cell electrical activity and Ca^2+^ handling. For example, one study determined that pharmacological GIRK channel inhibition blocks adrenaline-induced *V*_m_ hyperpolarization in rat β-cells (36), but another manuscript found that GIRK channel inhibition does not prevent adrenaline-induced *V*_m_ hyperpolarization in mouse β-cells (19). Furthermore, treatment with a wide range of other K^+^ channel blockers failed to inhibit adrenaline-induced β-cell *V*_m_ hyperpolarization. These observations suggest that G_i/o_ signaling-induced β-cell *V*_m_ hyperpolarization not mediated by GIRK or other K^+^ channels.

Electrogenic Na^+^/K^+^ ATPases (NKAs) can be activated by G_i/o_-coupled GPCR signaling leading to *V*_m_ hyperpolarization (38, 39). NKAs preserve steep ionic gradients that are essential for setting and maintaining β-cell *V*_m_ by extruding three intracellular Na^+^ ions in exchange for two extracellular K^+^ ions; this net outward cationic flux hyperpolarizes *V*_m_ (35, 38, 39). The NKA α1 subunit is highly expressed in mouse and human β-cells, showing greater than 100x the transcript expression of the most abundant GIRK channel transcript (e.g. *KCNJ6)* (11–13). Inhibition of islet NKAs with ouabain leads to β-cell *V*_m_ depolarization and enhances insulin secretion (40, 41). Ouabain also blocks glucose-stimulated [Ca^2+^]_i_ oscillations and decreases insulin receptor-mediated *V*_m_ hyperpolarization in K_ATP_ deficient SUR1^-/-^ islets (35). Furthermore, glucose inhibits β-cell NKA activity through protein kinase C (PKC) and phospholipase A2 dependent pathways, which promotes β-cell glucose-stimulated Ca^2+^ influx and presumably GSIS (42). Moreover, G_i/o_ signaling controls the activity of several β-cell protein kinases known to regulate NKA function. For example, protein kinase A (PKA), which limits NKA activity, is inhibited by SSTR signaling in a cAMP-dependent manner; whereas, sarcoma (Src) tyrosine kinases augment NKA function and are activated by SSTR signaling (38, 43–47). Thus, G_i/o_-coupled GPCRs are predicted to influence β-cell Ca^2+^ handling through changes in NKA α1 subunit phosphorylation.

Here we show for the first time, to the best of our knowledge, that NKA-mediated β-cell *V*_m_ hyperpolarization is a conserved mechanism for G_i/o_-coupled GPCR control of β-cell Ca^2+^ handling and insulin secretion. Activation of G_i/o_ signaling in β-cells generated ouabain- and K^+^-sensitive outward currents as well as [Ca^2+^]_i_ oscillations independently of K_ATP_ in both mouse and human β-cells. Inhibition of SST secretion from δ-cells increased islet [Ca^2+^]_i_, accelerated islet [Ca^2+^]_i_ oscillations, and enhanced GSIS, demonstrating the critical role of δ-cell paracrine signaling in tuning β-cell Ca^2+^ handling and insulin secretion, likely through NKA activation. β-cell NKA function also decreased in a cAMP-dependent manner due to activation of PKA. Furthermore, stimulation of tyrosine kinase activity (Src or insulin receptors) initiated islet [Ca^2+^]_i_ oscillations, which may indicate that tyrosine kinase signaling serves as a crucial mechanism for regulation of β-cell NKAs. Therefore, these findings illuminate a conserved mechanism for G_i/o_-coupled GPCR control of β-cell NKAs that serves a key role in regulating β-cell electrical Excitability, Ca^2+^ handling, and insulin secretion.

## RESULTS

### SSTR signaling activates GIRK channel-independent outward currents

It is generally accepted that β-cell G_i/o_ signaling activates hyperpolarizing GIRK channels (16, 36); however, measurement of β-cell GIRK currents has proven difficult utilizing traditional voltage-clamp recording techniques. Thus, we employed a modified recording paradigm to elicit quantifiable SST-induced β-cell currents. SST-mediated changes in β-cell *V*_m_ were monitored in intact islets and whole-cell currents were recorded in response to voltage ramps before and after each treatment. In C57 control islets stimulated with 20mM glucose and 1mM tolbutamide, SST induced outward currents with little rectification (current amplitude at −50mV: 15.7±1.9pA; Fig. 1A–1C; *P*<0.0001); SST also hyperpolarized *V*_m_ (−30.2±3.5mV; Fig. 1A and 1D; *P*<0.0001). As transcriptome studies indicate that *Kcnj6* is the most abundant GIRK channel transcript (11–13), GIRK2 deficient (GIRK2 KO^Panc^) islets were utilized to determine the contribution of GIRK2 channels to SST-induced β-cell currents. SST treatment elicited similar outward non-rectifying currents in β-cells without GIRK2 channels (current amplitude at −50mV: 17.6±2.0pA; Fig. 1E and 1F; *P*<0.0001) and hyperpolarized *V*_m_ (−23.2±2.6mV; Fig. 1G; *P*<0.01). Moreover, SST triggered [Ca^2+^]_i_ oscillations in 95.3±2.4% of islets without GIRK2 channels, which was indistinguishable from control islets (94.7±4.8%; Fig. 1H and 1I), and reduced [Ca^2+^]_i_ plateau fraction by 32.6±4.2% compared to before treatment (Fig. 1H and 1J; *P*<0.01). GIRK channels were also pharmacologically inhibited with 200nM tertiapin-Q to confirm GIRK channel-independent SST-induced β-cell currents. In the presence of tertiapin-Q, SST elicited outward currents with minimal inward rectification in control β-cells (max at −50mV: 12.4±1.8pA; Fig. 1K–1M; *P*<0.0001) and hyperpolarized *V*_m_ (−25.2±6.7mV; Fig. 1N; *P*<0.001). Furthermore, 76.8±10.5% of control islets treated with tertiapin-Q subsequent to SST continued to oscillate, which was not significantly different than with SST alone (92.8±7.3%; Fig. 1O and 1P). However, tertiapin-Q treatment after SST did modestly increase islet [Ca^2+^]_i_ plateau fraction by 13.7±4.8% (Fig. 1O and 1Q; *P*<0.05). These data demonstrate that SST-induced outward β-cell currents, which hyperpolarize *V*_m_ are largely independent of GIRK channel activity.

**Fig. 1:**
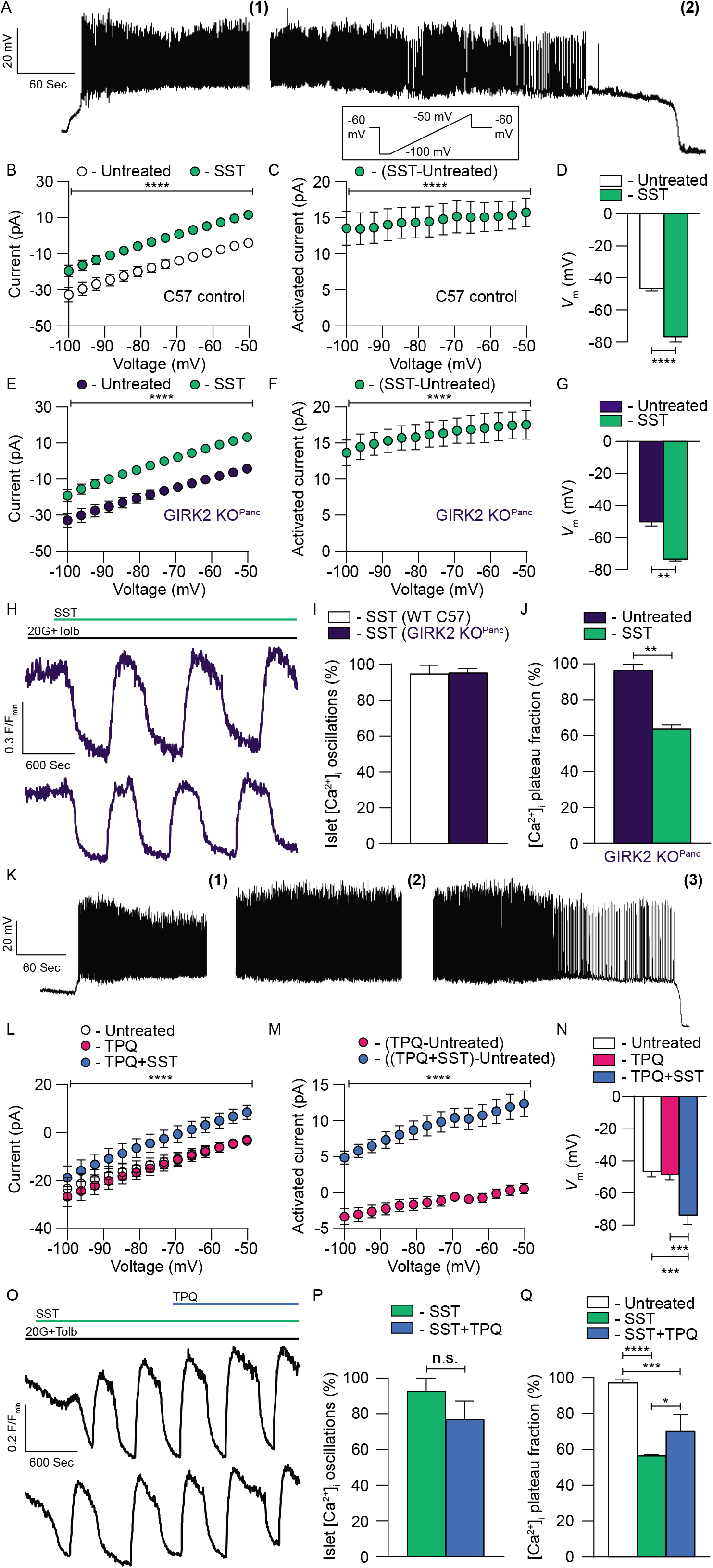
SSTR-mediated β-cell currents are not due to GIRK channel activation. (A) Representative β-cell *V*_m_ recording from an intact mouse islet. Whole-cell β-cell currents were measured in response to a voltage ramp protocol (inset) after treatment with **(1)** 20mM glucose (20G)+1mM tolbutamide (Tolb) and **(2)** 20G+1mM Tolb+200nM SST. (B) Average C57 β-cell currents with 20G+1mM Tolb before (white) and after SST (green; *n*=11 β-cells in intact islets; \*\*\*\**P*<0.0001). (C) Average SST-induced C57 β-cell currents (green; *n*=11 β-cells in intact islets; \*\*\*\**P*<0.0001). (D) Average C57 β-cell *V*_m_ with 20G+Tolb before (white) and after SST (green; *n*=13 β-cells in intact islets; \*\*\*\**P*<0.0001). (E) Average GIRK2 KO^Panc^ β-cell currents with 20G+Tolb before (dark blue) and after SST (green; *n*=4 β-cells in intact islets; \*\*\*\**P*<0.0001). (F) Average SST-induced GIRK2 KO^Panc^ β-cell currents (green; *n*=4 β-cells in intact islets; \*\*\*\**P*<0.0001). (G) Average GIRK2 KO^Panc^ β-cell *V*_m_ with 20G+Tolb before (dark blue) and after SST (green; *n*=4 β-cells in intact islets; \*\**P*<0.01). (H) Representative normalized (F/F_min_) C57 islet Fura-2 Ca^2+^ responses to 200nM SST in the presence of 20G+1mM Tolb. (I) Average percentage of C57 (white; *n*=islets from 6 mice) and GIRK2 KO^Panc^ (dark blue; *n*=islets from 3 mice) islets displaying SST-induced [Ca^2+^]_i_ oscillations. (J) Average GIRK2 KO^Panc^ islet [Ca^2+^]_i_ plateau fraction (defined as Fura-2 fluorescence after treatment≥50% of average Fura-2 fluorescence before SST treatment) with 20G+Tolb before (dark blue) and after SST (green; *n*=islets from 3 mice; \*\**P*<0.01). (K) Representative β-cell *V*_m_ recording from an intact mouse islet. Whole-cell β-cell currents were measured in response to a voltage ramp protocol after treatment with **(1)** 20G+1mM Tolb, **(2)** 20G+1mM Tolb+200nM tertiapin-Q (TPQ), and **(3)** 20G+1mM Tolb+200nM TPQ+200nM SST. (L) Average C57 β-cell currents with 20G+Tolb (white), with TPQ (pink), and with TPQ+SST (blue; *n*=8 β-cells in intact islets; \*\*\*\**P*<0.0001). (M) Average TPQ-induced (pink) and TPQ+SST-induced C57 β-cell currents (green; *n*=8 β-cells in intact islets; \*\*\*\**P*<0.0001). (N) Average C57 β-cell *V*_m_ with 20G+Tolb (white), after TPQ (pink), and after TPQ+SST (blue; *n*=8 β-cells in intact islets; \*\*\**P*<0.001). (O) Representative normalized (F/F_min_) C57 islet Fura-2 Ca^2+^ responses to 200nM SST and 200nM TPQ in the presence of 20G+1mM Tolb. (P) Average percentage of C57 islets with 20G+Tolb displaying [Ca^2+^]_i_ oscillations in response to SST (green) and SST+TPQ (blue; *n*=islets from 4 mice). (Q) Average C57 islet [Ca^2+^]_i_ plateau fraction with 20G+Tolb (white), after SST (green), and after SST+TPQ (blue; *n*=islets from 4 mice; \**P*<0.05, \*\*\**P*<0.001, and ****P<0.0001). Statistical analysis was conducted using paired two-sample t-tests (B-G, L, and M), unpaired two-sample t-tests (I, J, and P), or one-way ANOVA (N and Q); uncertainty is expressed as mean ± SEM.

### SSTR signaling induces islet [Ca^2+^]_i_ oscillations by stimulating β-cell NKA activity

SST stimulated outward currents below the equilibrium potential of K^+^, which indicates that SST-induced β-cell currents are not mediated by K^+^ channels (Fig. 1C, 1F, and 1M). This suggests that SST-induced outward currents are due to efflux of another cation such as Na^+^, which requires movement against the ion concentration gradient through energy-dependent ion pumps. Therefore, we investigated whether G_i/o_ signaling decreases β-cell [Ca^2+^]_i_ by facilitating Na^+^ efflux through electrogenic NKAs. SST-induced [Ca^2+^]_i_ oscillations stopped in 97.1 ±2.9% of islets following NKA inhibition with 150μM ouabain (Fig. 2A–2C; *P*<0.001). SST treatment also significantly decreased islet [Ca^2+^]_i_ plateau fraction with (54.8±5.8% decrease; Fig. 2D; *P*<0.0001) and without K_ATP_ inhibition (76.4±6.4% decrease; Fig. 2D; *P*<0.0001). Importantly, K_ATP_ activation with 125μM diazoxide during NKA inhibition decreased β-cell [Ca^2+^]_i_ by 82.5±15.7% (Fig. 2E and Fig. S1; *P*<0.01), which reinforces that SST-induced [Ca^2+^]_i_ oscillations are not due to activation of a K^+^ conductance. As NKA function is regulated by cAMP-dependent signaling pathways (38, 39), we also measured mouse islet [cAMP]_i_ along with [Ca^2+^]_i_ (Fig. 2A and 2B). Cross-correlation analysis of SST-mediated islet [Ca^2+^]_i_ and [cAMP]_i_ oscillations revealed a strong negative correlation between the two (Fig. 2F; max correlation coefficient: −0.46±0.02) with [Ca^2+^]_i_ oscillations preceding [cAMP]_i_ oscillations by approximately 100ms. However, [cAMP]_i_ only modestly decreased following SST treatment indicating that changes in [cAMP]_i_ are not responsible for SST activation of β-cell NKAs. The importance of SST-induced NKA activity to β-cell Ca^2+^ handling was confirmed by removing extracellular K^+^, which is required for NKA function. Following removal of extracellular K^+^ SST-induced [Ca^2+^]_i_ oscillations ceased in 96.1 ±1.9% of islets (Fig. 1C and 1G; *P*<0.0001); K^+^ supplementation rapidly restored [Ca^2+^]_i_ oscillations (Fig. 1F). Finally, we confirmed that SST does not induce β-cell currents or hyperpolarize *V*_m_ in the absence of extracellular K^+^ utilizing patch-clamp electrophysiology (Fig. 1H–1K). Taken together, these findings establish that SST-induced NKA activation decreases β-cell [Ca^2+^]_i_. Moreover, our results show that G_i/o_-coupled GPCR signaling induces oscillations in both [Ca^2+^]_i_ and [cAMP]_i_, which likely results from oscillations in NKA activity.

**Fig. 2:**
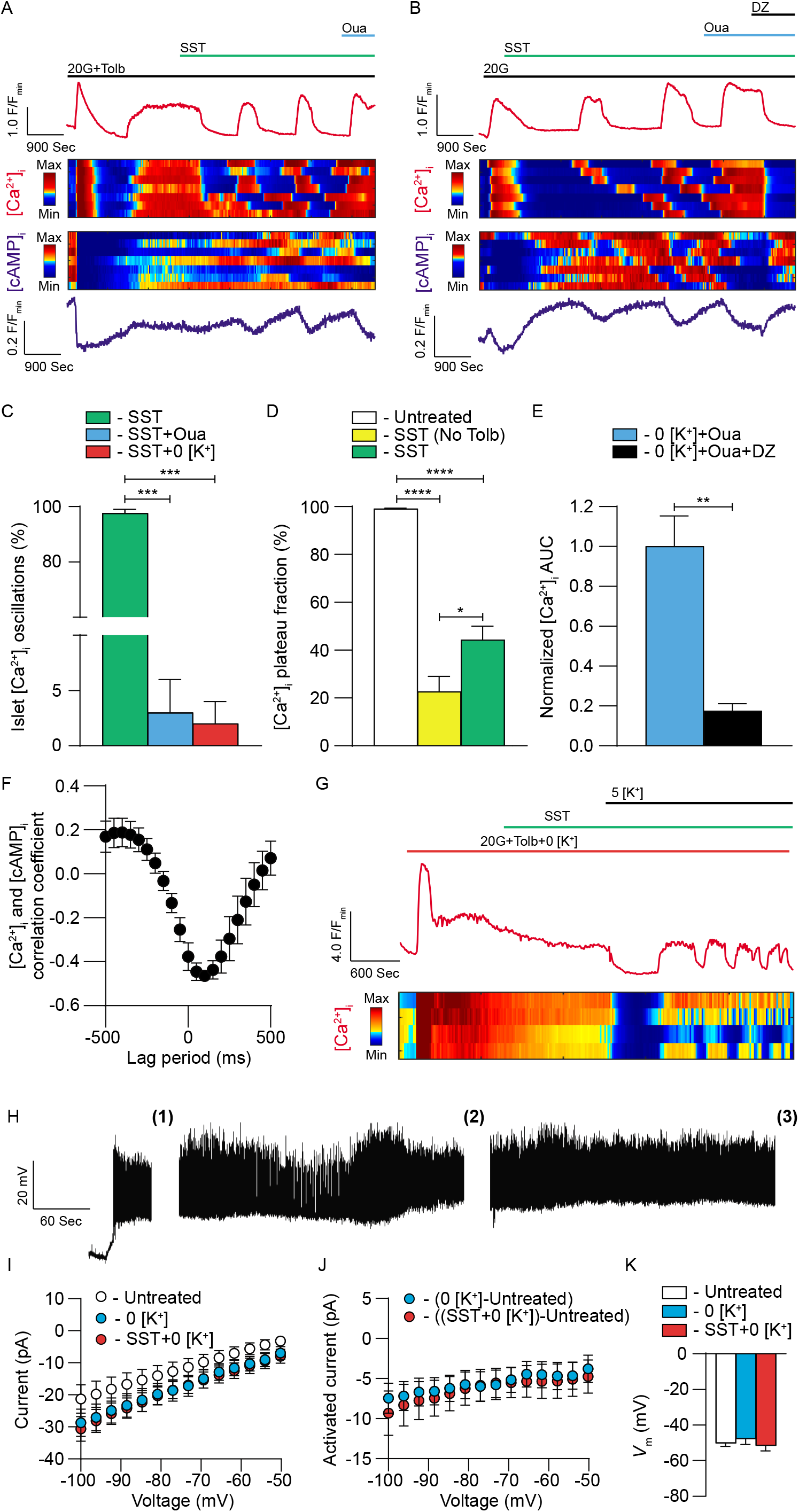
SST-induced activation of β-cell NKAs generates islet [Ca^2+^]_i_ and [cAMP]_i_ oscillations. (A) Representative C57 islet jRGECO1a [Ca^2+^]_i_ (top, red) and cAMPr [cAMP]_i_ (bottom, blue) responses to 200nM SST and 150μM ouabain (Oua) in the presence of 20G+1mM Tolb. Heatmaps illustrating typical islet [Ca^2+^]_i_ (middle, upper) and [cAMP]_i_ (middle, lower) responses from a single experiment. (B) Representative C57 islet jRGECO1a [Ca^2+^]_i_ and cAMPr [cAMP]_i_ responses to 200nM SST, 150μM Oua, and 125μM diazoxide (DZ) in the presence of 20G. Heatmaps illustrating typical islet [Ca^2+^]_i_ and [cAMP]_i_ responses from a single experiment. (C) Average percentage of C57 islets with 20G+1mM Tolb displaying [Ca^2+^]_i_ oscillations in response to SST (green; *n*=islets from 7 mice), SST+Oua (light blue; *n*=islets from 3 mice), and SST+0mM K^+^ (0 [K^+^]; orange; *n*=islets from 4 mice; \*\*\**P*<0.001). (D) Average C57 islet [Ca^2+^]_i_ plateau fraction with 20G+Tolb (white; *n*=islets from 10 mice), with 20G+SST (yellow; *n*=islets from 6 mice), and with 20G+Tolb+SST (green; *n*=islets from 4 mice; \**P*<0.05 and ****P<0.0001). (E) Average C57 islet [Ca^2+^]_i_ area under the curve (AUC; averaged over at least 10 minutes) normalized to 20G+0 [K^+^] before (light blue; *n*=islets from 4 mice) and after DZ (black; *n*=islets from 4 mice; \*\**P*<0.01). (F) Cross-correlation analysis of C57 islet [Ca^2+^]_i_ and [cAMP]_i_ (*n*=islets from 4 mice). (G) Representative C57 islet jRGECO1a [Ca^2+^]_i_ (top, red) response to 200nM SST and 5mM K^+^ (5 [K^+^]) in the presence of 20G+0 [K^+^]. Heatmap illustrating typical islet [Ca^2+^]_i_ responses (bottom) from a single experiment. (H) Representative β-cell *V*_m_ recording from an intact mouse islet. Whole-cell β-cell currents were measured in response to a voltage ramp protocol after treatment with **(1)** 20G+1mM Tolb, **(2)** 20G+1mM Tolb+0 [K^+^], and **(3)** 20G+1mM Tolb+0 [K^+^]+200nM SST. (I) Average C57 β-cell currents with 20G+1mM Tolb (white), after 0 [K^+^] (light blue), and after 0 [K^+^]+SST (orange; *n*=10 β-cells in intact islets). (J) Average 0 [K^+^]-induced (light blue) and 0 [K^+^]+SST-induced C57 β-cell currents (orange; *n*=10 β-cells in intact islets). (K) Average C57 β-cell *V*_m_ with 20G+Tolb (white; *n*=10 β-cells in intact islets), after 0 [K^+^] (light blue; 9 β-cells in intact islets), and after 0 [K^+^]+SST (orange; *n*=6 β-cells in intact islets). Statistical analysis was conducted using paired two-sample t-tests (I and J), unpaired two-sample t-tests (E), or one-way ANOVA (C, D, and K); uncertainty is expressed as mean ± SEM.

NKA activity would be predicted to maintain low β-cell [Na^+^]_i_, therefore, we simultaneously measured islet [Na^+^]_i_ along with [Ca^2+^]_i_. Cross-correlation analysis of SST-induced islet [Na^+^]_i_ and [Ca^2+^]_i_ oscillations in the presence of 20mM glucose and 1mM tolbutamide demonstrated that changes in islet [Na^+^]_i_ closely follow [Ca^2+^]_i_ (Fig. 3A and 3B; max correlation coefficient: 0.67±0.8). Interestingly, there was also a strong cross-correlation between islet [Na^+^]_i_ and [Ca^2+^]_i_ during glucose-stimulated [Ca^2+^]_i_ oscillations at 9mM glucose (Fig. 3C and 3D; max correlation coefficient: 0.75±0.11), which suggests that NKA regulation of β-cell Ca^2+^ handling is not restricted to G_i/o_ signaling. Moreover, as islet [Na^+^]_i_ oscillates in response to SST, this indicates that SSTR regulation of β-cell NKA function is also oscillatory.

**Fig. 3:**
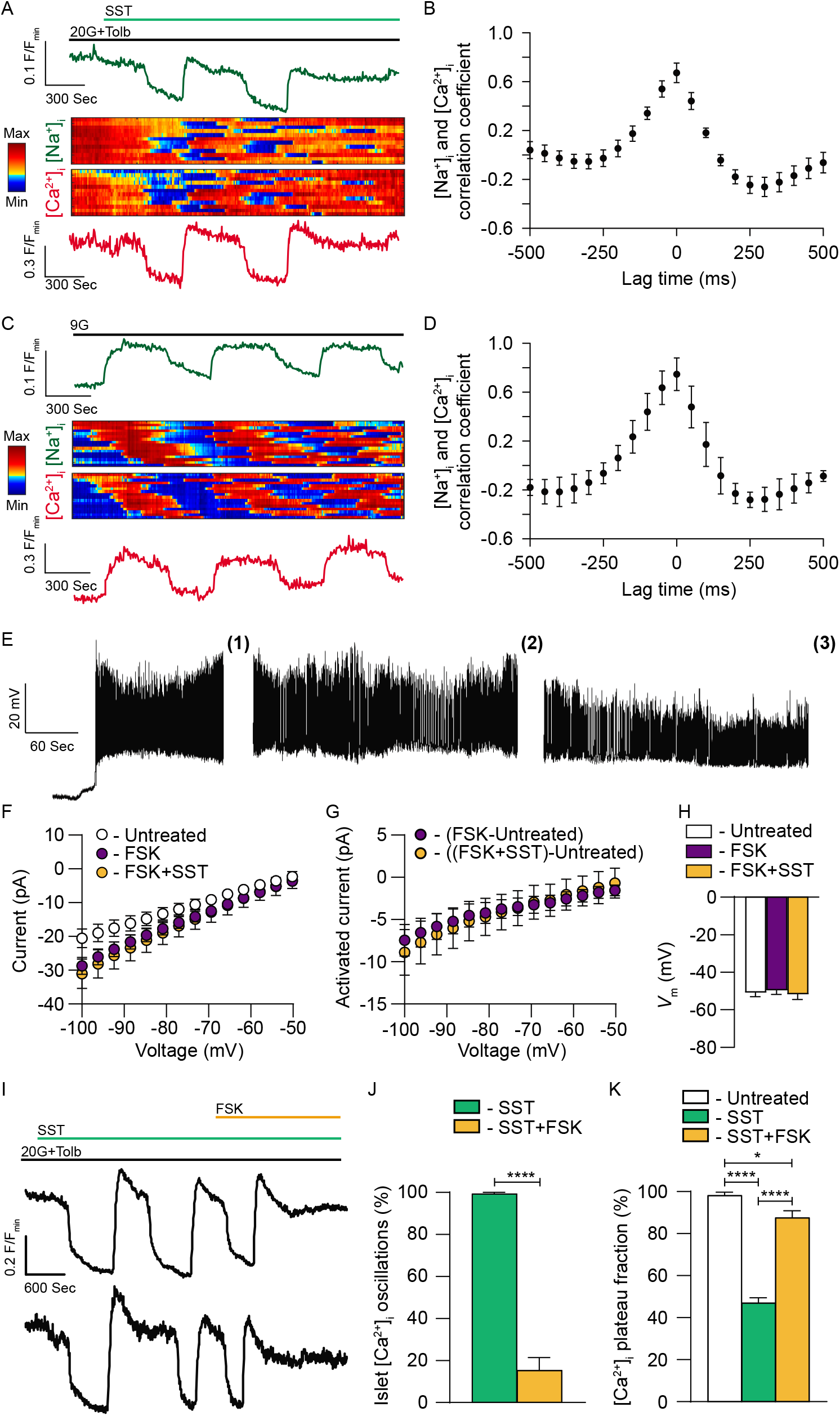
NKA-mediated islet [Ca^2+^]_i_ oscillations are inhibited by forskolin-induced increases in [cAMP]_i_. (A) Representative C57 islet ING-2 [Na^+^]_i_ (top, dark green) and Fura Red AM [Ca^2+^]_i_ (bottom, red) responses to 200nM SST in the presence of 20G+1mM Tolb. Heatmaps illustrating typical islet [Na^+^]_i_ (middle, upper) and [Ca^2+^]_i_ (middle, lower) responses from a single experiment. (B) Cross-correlation analysis of C57 islet [Na^+^]_i_ and [Ca^2+^]_i_ with 20G+Tolb+SST (*n*=islets from 4 mice). (C) Representative C57 islet ING-2 [Na^+^]_i_ and Fura Red [Ca^2+^]_i_ responses to 9mM G (9G). Heatmaps illustrating typical islet [Na^+^]_i_ and [Ca^2+^]_i_ responses from a single experiment. (D) Cross-correlation analysis of C57 islet [Na^+^]_i_ and [Ca^2+^]_i_ with 9G (*n*=islets from 3 mice). Representative β-cell *V*_m_ recording from an intact mouse islet. Whole-cell β-cell currents were measured in response to a voltage ramp protocol after treatment with **(1)** 20G+1mM Tolb, **(2)** 20G+1mM Tolb+5μM forskolin (FSK), and **(3)** 20G+1mM Tolb+5μM FSK+200nM SST. (F) Average C57 β-cell currents with 20G+1mM Tolb (white), after FSK (purple), and after FSK+SST (light orange; *n*=11 β-cells in intact islets). (G) Average FSK-induced (purple) and FSK+SST-induced C57 β-cell currents (light orange; *n*=11 β-cells in intact islets). (H) Average C57 β-cell *V*_m_ with 20G+Tolb (white), after FSK (purple), and after FSK+SST (light orange; *n*=11 β-cells in intact islets). (I) Representative normalized (F/F_min_) C57 islet Fura-2 Ca^2+^ responses to 200nM SST and 5μM FSK in the presence of 20G+1mM Tolb. (J) Average percentage of C57 islets with 20G+Tolb displaying [Ca^2+^]_i_ oscillations in response to SST (green) and SST+FSK (light orange; *n*=islets from 4 mice). (K) Average C57 islet [Ca^2+^]_i_ plateau fraction with 20G+Tolb before (white), after SST (green), and after SST+FSK (light orange; *n*=islets from 4 mice; \**P*<0.05 and \*\*\*\**P*<0.0001). Statistical analysis was conducted using paired two-sample t-tests (F and G), unpaired two-sample t-tests (J), or one-way ANOVA (H and K); uncertainty is expressed as mean ± SEM.

It is established that elevations in [cAMP]_i_ inhibit NKA activity and we observed a correlation between increasing [cAMP]_i_ and termination of SST-induced islet [Ca^2+^]_i_ oscillations. Thus, we utilized forskolin to increase islet [cAMP]_i_ and measured changes in SST-induced β-cell NKA activity. Treatment with 5μM forskolin blocked SST-induced β-cell NKA currents and prevented *V*_m_ hyperpolarization (Fig. 3E–3H). Furthermore, SST-induced [Ca^2+^]_i_ oscillations persisted in only 15.3±6.2% of islets following forskolin treatment compared with SST alone (84.8±6.2% decrease; Fig. 3I and 3J; *P*<0.0001); after forskolin treatment [Ca^2+^]_i_ plateau fraction also increased by 40.5±2.9% compared to with SST alone (Fig. 3I and 3K; *P*<0.0001). These results show that cAMP blocks SST-induced islet [Ca^2+^]_i_ oscillations by inhibiting β-cell NKA activity.

### NKA activation is a conserved mechanism for G_i/o_-coupled GPCR control of β-cell [Ca^2+^]_i_

β-cells express a number of G_i/o_-coupled GPCRs in addition to SSTRs including α2A ADRs and D2-like DRDs (11–15). Thus, we examined whether β-cell NKA activation is a conserved mechanism for G_i/o_ signaling-mediated control of islet [Ca^2+^]_i_. Islets stimulated with 20mM glucose and 1mM tolbutamide did not oscillate, whereas all islets treated with the α-ADR activator clonidine (200nM) exhibited [Ca^2+^]_i_ oscillations (Fig 4A and 4B; *P*<0.0001) and a 64.0±3.3% decrease in [Ca^2+^]_i_ plateau fraction (Fig. 4A–4C; *P*<0.0001). Treatment with ouabain terminated clonidine-induced [Ca^2+^]_i_ oscillations in 94.9±3.0% of islets (Fig. 4A and 4B; *P*<0.0001) and increased [Ca^2+^]_i_ plateau fraction by 61.1±3.4% (Fig. 4A and 4C; *P*<0.0001), which was indistinguishable from islets before clonidine treatment. These findings indicate that stimulation of NKA activity is a conserved mechanism for G_i/o_ -coupled GPCR control of β-cell [Ca^2+^]_i_.

**Fig. 4:**
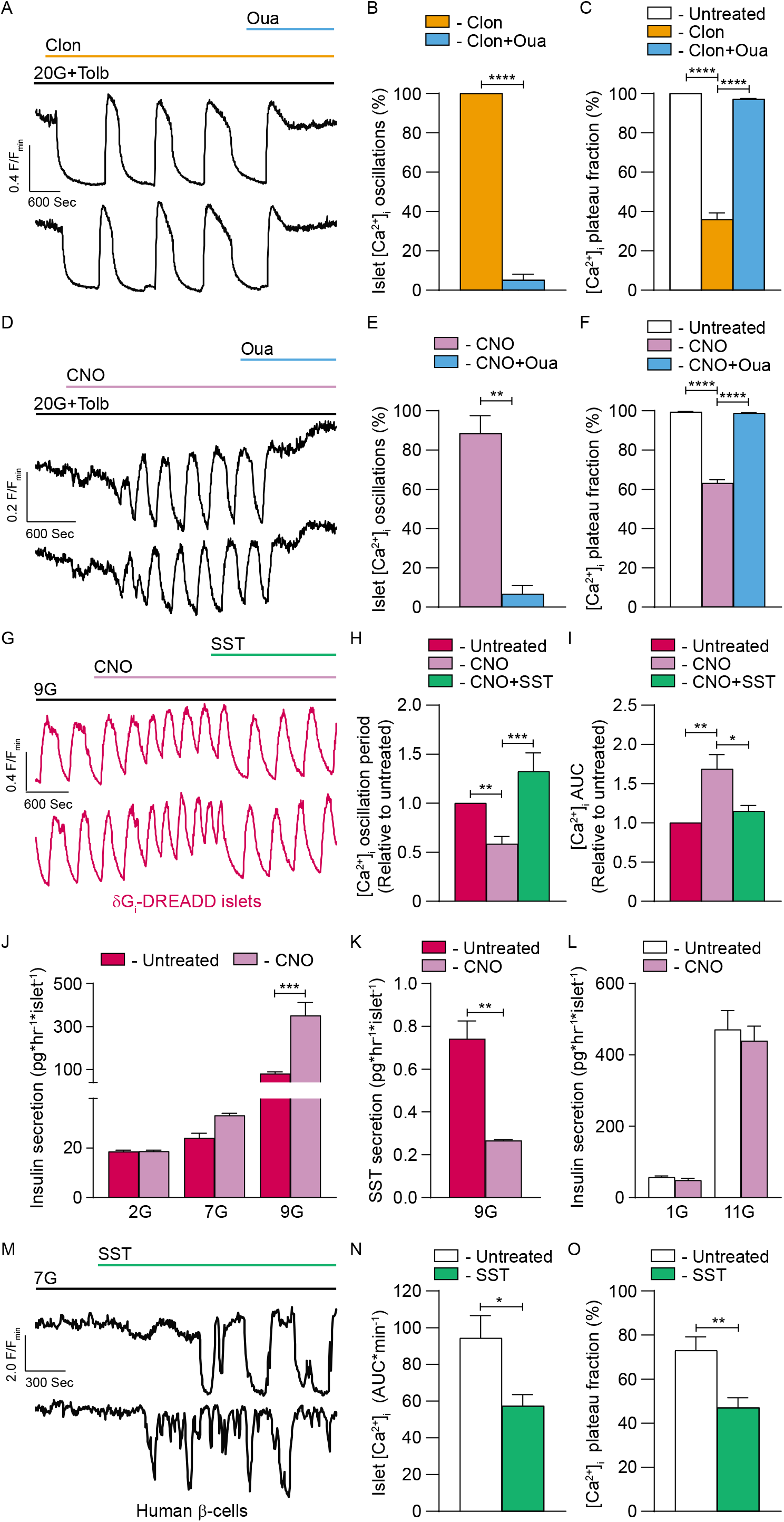
Activation of β-cell NKAs is a conserved mechanism for G_i/o_-coupled GPCR control of islet Ca^2+^ handling: (A) Representative normalized (F/F_min_) C57 islet Fura-2 Ca^2+^ responses to 200nM clonidine (Clon) and 150μM Oua in the presence of 20G+1mM Tolb. (B) Average percentage of C57 islets with 20G+Tolb displaying [Ca^2+^]_i_ oscillations in response to Clon (orange) and Clon+Oua (light blue; *n*=islets from 3 mice). (C) Average C57 islet [Ca^2+^]_i_ plateau fraction with 20G+Tolb (white), after Clon (orange), and after Clon+Oua (light blue; *n*=islets from 3 mice; \*\*\*\**P*<0.0001). (D) Representative normalized (F/F_min_) Fura-2 Ca^2+^ responses of C57 islets transduced with RIP-Gi DREADD LVs to 10μM CNO and 150μM Oua in the presence of 20G+1mM Tolb. (E) Average percentage of C57 islets transduced with RIP-Gi DREADD LVs with 20G+Tolb displaying [Ca^2+^]_i_ oscillations in response to CNO (light pink) and CNO+Oua (light blue; *n*=islets from 3 mice; \*\**P*<0.01). (F) Average C57 islet [Ca^2+^]_i_ plateau fraction transduced with RIP-Gi DREADD LVs with 20G+Tolb (white), after CNO (light pink), and after CNO+Oua (light blue; *n*=islets from 3 mice; \*\*\*\**P*<0.0001). (G) Representative normalized (F/F_min_) δGi DREADD islet Cal-590 Ca^2+^ responses to 10μM CNO and 200nM SST in the presence of 9mM glucose (9G). (H) Average δGi DREADD islet [Ca^2+^]_i_ oscillation period relative to before treatment (dark pink; *n*=islets from 7 mice) following CNO (light pink; *n*=islets from 7 mice) and CNO+SST (green; *n*=islets from 4 mice; \*\**P*<0.01 and \*\*\**P*<0.001). (I) Average δGi DREADD islet [Ca^2+^]_i_ AUC (sum of 15 minutes) relative to before treatment (dark pink), after CNO (light pink), and after CNO+SST (green; *n*=islets from 4 mice; \**P*<0.05 and \*\**P*<0.01). (J) Average insulin secretion from δGi DREADD islets without CNO (dark pink) and with CNO (light pink) at 2mM glucose (2G; *n*=islets from 3 mice), 7mM glucose (7G; *n*=islets from 3 mice), and 9G (*n*=islets from 6 mice; \*\*\**P*<0.001). (K) Average SST secretion from δGi DREADD islets without CNO (dark pink) and with CNO (light pink) at 9G (*n*=islets from 3 mice; \*\**P*<0.01). (L) Average insulin secr etion from C57 islets without CNO (white) and with CNO (light pink) at 1mM glucose (1G; *n*=islets from 3 mice) and at 11mM glucose (11G; *n*=islets from 3 mice). (M) Representative normalized (F/F_min_) human β-cell jRGECO1a [Ca^2+^]_i_ responses (within intact human islets) to 400nM SST in the presence of 7G. (N) Average human β-cell [Ca^2+^]_i_ AUC (averaged over at least 15 minutes) with 7G before (white) and after SST (green; *n*=islets from 4 healthy donors; \**P*<0.05). (O) Average human β-cell [Ca^2+^]_i_ plateau fraction (within intact human islets) with 7G before (white) and after SST (green; *n*=islets from 4 healthy donors; \*\**P*<0.01). Statistical analysis was conducted using unpaired two-sample t-tests (B, E, K, N, and O), or one-way ANOVA (C, F, H-J, and L); uncertainty is expressed as mean ± SEM.

SSTRs and ADRs have also been shown to control α-cell function (16, 48, 49), which is predicted to impact β-cell Ca^2+^ handling. Therefore, G_i/o_-coupled DREADDS driven by an optimized RIP were employed to selectively activate G_i/o_ signaling in β-cells (Fig. 4D). Islets displayed no [Ca^2+^]_i_ oscillations in the presence of 20mM glucose and 1mM tolbutamide; however, following treatment with 10μM CNO [Ca^2+^]_i_ oscillations were observed in 88.5±9.0% of islets (Fig. 4D and 4E) and [Ca^2+^]_i_ plateau fraction decreased 36.3±% (Fig. 4D and 4F; *P*<0.0001). Interestingly, only a small subset of β-cells expressed G_i/o_-coupled DREADDS in each islet (based on mCherry fluorescence), suggesting that electrical coupling between β-cells amplifies G_i/o_ signaling-induced NKA activation. Following ouabain treatment, [Ca^2+^]_i_ oscillations ceased in 93.4±4.3% of islets (Fig. 4D and 4E; *P*<0.01) and [Ca^2+^]_i_ plateau fraction increased 35.7±1.8% (Fig. 4D and 4F, *P*<0.0001), which was indistinguishable from islets before CNO treatment. These data confirm that direct stimulation of β-cell G_i/o_ signaling initiates islet [Ca^2+^]_i_ oscillations.

We next sought to determine if endogenous islet SST controls [Ca^2+^]_i_ oscillations; this was accomplished utilizing islets from mice selectively expressing G_i/o_-coupled DREADDs in δ-cells (δGi-DREADDs) to inhibit SST secretion during glucose-stimulated [Ca^2+^]_i_ oscillations. Activation of δ-cell G_i/o_ signaling with CNO under stimulatory (9mM) glucose conditions decreased the period of [Ca^2+^]_i_ oscillations by 41.7±7.7% (Fig. 4G and 4H; *P*<0.01) and increased [Ca^2+^]_i_ area under the curve (AUC) by 68.8±18.3% (Fig. 4G and 4I; *P*<0.01) relative to before CNO treatment. This effect was reversed following treatment with exogenous SST (Fig. 4G–4I); [Ca^2+^]_i_ oscillation period increased by 74.1±20.4% (Fig. 4G and 4H: *P*<0.001) and [Ca^2+^]_i_ AUC decreased by 54.0±19.8% (Fig. 4G and 4I; *P*<0.05). As stimulation of δ-cell G_i/o_ signaling enhanced β-cell Ca^2+^ influx and accelerated [Ca^2+^]_i_ oscillation frequency, we examined the effect of δ-cell G_i/o_-coupled GPCR activation on SST and insulin secretion. CNO activation of δGi-DREADDs had no effect on insulin secretion under low (2mM) or basal (7mM) glucose conditions; however, under stimulatory (9mM) glucose conditions δ-cell G_i/o_ signaling increased insulin secretion from 81.5±7.7 to 352.2±59.4pg insulin·hr^-1^·islet^-1^ (Fig. 4J; *P*<0.0001). Importantly, under identical conditions δGi-DREADD activation decreased SST secretion from 0.743±0.082 to 0.267±0.003pg SST·hr^-1^islet^-1^ (Fig. 4K; *P*<0.01). Insulin secretion from control islets was not affected by CNO at either low (1mM) or stimulatory (11mM) glucose conditions (Fig. 4L). These findings show that δ-cell SST secretion regulates islet Ca^2+^ handling and insulin secretion under physiological conditions. The data also suggest that SSTR-mediated activation of β-cell NKAs decelerates glucose-stimulated [Ca^2+^]_i_ oscillations and resulting pulsatile insulin secretion.

### G_i/o_-coupled GPCRs control human islet Ca^2+^ handling by increasing β-cell NKA activity

As SSTR signaling hyperpolarizes human β-cell *V*_m_ and inhibits voltage-dependent Ca^2+^ channel activity (16), we went on to examine the mechanisms responsible. A GCaMP6s genetically encoded [Ca^2+^]_i_ indicator driven by an optimized RIP was utilized to examine SSTR control of human β-cell [Ca^2+^]_i_ (50). Treatment with exogenous SST (400nM) during stimulatory (7mM) glucose conditions decreased β-cell [Ca^2+^]_i_ AUC by 37.2±0.8% (Fig. 4M and 4N; *P*<0.05) and [Ca^2+^]_i_ plateau fraction by 36.4±2.9% (Fig. 4M and 4O; *P*<0.01). These results strongly indicate that G_i/o_ signaling regulates β-cell [Ca^2+^]_i_ in human islets.

Transcriptional analysis shows that human β-cells express high levels of *ATP1A1* transcript (gene encoding the NKA α1 subunit) (11–13). Indeed, immunofluorescence staining of human pancreatic sections confirmed that insulin positive β-cells stain positive for NKA α1, and revealed that the protein is predominantly restricted to cell membranes (Fig. 5A). Interestingly, other islet cells stained positive for NKA α1, which may indicate that NKA serves additional roles in pancreatic α- and/or δ-cells.

**Fig. 5:**
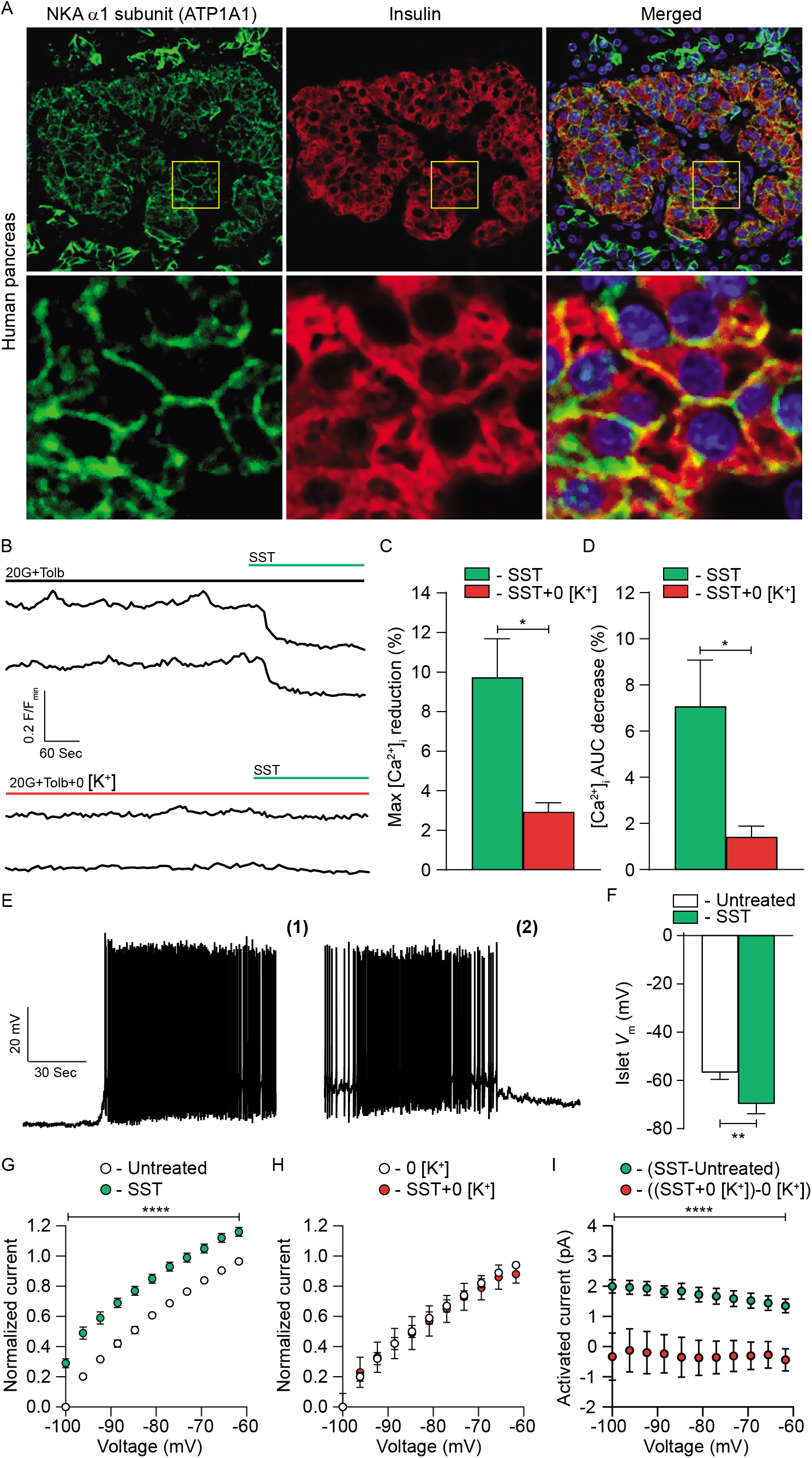
G_i/o_-coupled GPCRs regulate human β-cell electrical excitability by stimulating NKA activity: (A) Top row: Representative immunofluorescent staining of healthy human pancreatic sections for NKA α1 subunits (green), insulin (red), and a merged image of the two showing colocalization (yellow). Bottom row: Magnification of the corresponding areas outlined with yellow boxes above. (B) Representative human islet Cal-590 Ca^2+^ responses to 400nM SST in the presence of 20G+1mM Tolb with (top) and without extracellular K^+^ (0 [K^+^]; bottom). (C) Average maximum SST-induced decrease in human islet [Ca^2+^]_i_ relative to before treatment with (green) and without extracellular K^+^ (red; *n*=human islets from 5 healthy donors; \**P*<0.05). (D) Average SST-induced decrease in human islet [Ca^2+^]_i_ AUC (sum of 5 minutes) relative to before treatment with (green) and without extracellular K^+^ (red; *n*=human islets from 5 healthy donors; \**P*<0.05). (E) Representative β-cell *V*_m_ recording from an intact human islet. Whole-cell human β-cell currents were measured in response to a voltage ramp protocol after treatment with **(1)** 20G+1mM Tolb and **(2)** 20G+1mM Tolb+400nM SST. (F) Average human β-cell *V*_m_ with 20G+Tolb before (white) and after SST (green; *n*=7 β-cells in intact human islets; \*\**P*<0.01). (G) Average normalized (I/Imin) human β-cell currents with 20G+1mM Tolb before (white) and after SST (green; *n*=8 β-cells in intact human islets; \*\*\*\**P*<0.0001); Imin is defined as the minimum current measured before SST. (H) Average normalized (I/Imin) human β-cell currents with 20G+1mM Tolb+0 [K^+^] before (white) and after SST (red; *n*=5 β-cells in intact human islets). (I) Average SST-induced human β-cell currents with (green; *n*=8 β-cells in intact human islets; \*\*\*\**P*<0.0001) and without extracellular K^+^ (red; *n*=5 β-cells in intact islets). Statistical analysis was conducted using unpaired two-sample t-tests (C and D) or paired two-sample t-tests (F-I); uncertainty is expressed as mean ± SEM.

We next set out to determine whether G_i/o_-coupled GPCR signaling controls human β-cell [Ca^2+^]_i_ by augmenting NKA function. In human islets stimulated with 20mM glucose and 1mM tolbutamide, SST transiently decreased human islet [Ca^2+^]_i_ by 9.7±1.9% and reduced [Ca^2+^]_i_ AUC by 7.1±2.0% (Fig. 5B–5D). The effect of SST was greatly attenuated when extracellular K^+^ was excluded; SST-induced decreases in islet [Ca^2+^]_i_ were 3.6±0.3 fold lower and reductions in islet [Ca^2+^]_i_ AUC were 5.1±0.4 fold lower (Fig. 5B–5D; *P*<0.05). SST also hyperpolarized β-cell *V*_m_ from −56.8±2.8 to −69.7±4.1mV (*P*<0.01) and activated outward currents (current amplitude at −100 mV: 2.0±0.2pA; Fig. 5E–5I; *P*<0.0001). Importantly, SST-induced human β-cell currents were outward at voltages below the equilibrium potential of K^+^, which suggests Na^+^ movement through NKAs. Indeed, SST-induced β-cell currents disappeared when K^+^ was removed from the bath solution. Taken together, these data indicate that SSTR signaling, and presumably G_i/o_ signaling in general, increases human β-cell NKA activity resulting in *V*_m_ hyperpolarization and decreased [Ca^2+^]_i_.

### β-cell NKA activity is regulated by PKA and tyrosine kinase signaling

Forskolin-mediated elevations in [cAMP]_i_ influence β-cell function by stimulating PKA signaling (51, 52), which has been shown to decrease NKA activity (38, 39, 46); thus, PKA was pharmacologically blocked with H89 to investigate its role in G_i/o_-coupled GPCR control of β-cell electrical activity and Ca^2+^ handling. Treatment with 10μM H89 terminated SST-induced [Ca^2+^]_i_ oscillations (Fig. 6A), reduced islet [Ca^2+^]_i_ AUC by 84.7±3.6% (Fig. 6A and 6B; *P*<0.001), and decreased [Ca^2+^]_i_ plateau fraction by 99.5±0.3% (Fig. 6A and 6C; *P*<0.0001) compared to SST alone. Furthermore, H89 abolished forskolin-mediated increases in islet [Ca^2+^]_i_ AUC (Fig. 6A and 6B) as well as [Ca^2+^]_i_ plateau fraction (Fig. 6A and 6C). These data demonstrate that PKA serves as a negative regulator of NKA function in β-cells; however, as [cAMP]_i_ decreased only modestly in response to SST it is unlikely that a reduction in PKA activity accounts for initiation of SST-induced islet [Ca^2+^]_i_ oscillations.

**Fig. 6:**
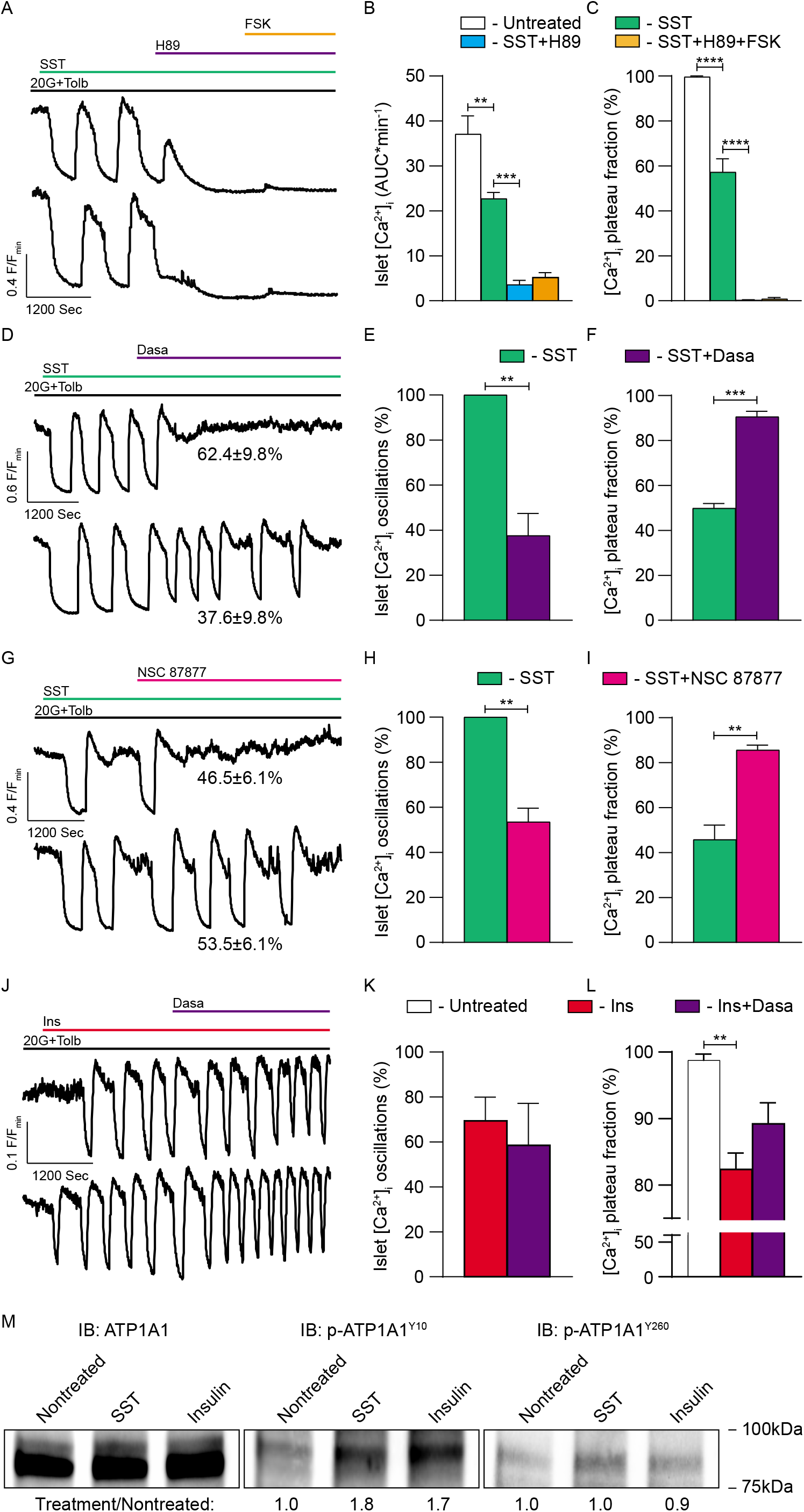
G_i/o_-coupled GPCR-mediated activation of β-cell NKAs is controlled by PKA and tyrosine kinase signaling: (A) Representative normalized (F/F_min_) C57 islet Fura-2 Ca^2+^ responses to 200nM SST, 10μM H89, and 5μM FSK in the presence of 20G+1mM Tolb. (B) Average C57 islet [Ca^2+^]_i_ AUC (averaged over at least 5 minutes) with 20G+Tolb (white), SST (green), SST+H89 (light blue), and SST+H89+FSK (light orange; *n*=islets from 3 mice; \*\**P*<0.01 and \*\*\**P*<0.001). (C) Average C57 islet [Ca^2+^]_i_ plateau fraction with 20G+Tolb (white), after SST (green), after SST+H89 (light blue), and after SST+H89+FSK (light orange; *n*=islets from 3 mice; \*\*\*\**P*<0.0001). (D) Representative normalized (F/F_min_) C57 islet Fura-2 Ca^2+^ responses to 200nM SST and 100nM dasatinib (Dasa) in the presence of 20G+1mM Tolb. Insets indicate the percentage of C57 islets exhibiting each type of Dasa [Ca^2+^]_i_ response. (E) Average percentage of C57 islets with 20G+Tolb displaying [Ca^2+^]_i_ oscillations in response to SST (green) and SST+Dasa (purple; *n*=islets from 3 mice; \*\**P*<0.01). (F) Average C57 islet [Ca^2+^]_i_ plateau fraction with 20G+Tolb after SST (green) and after SST+Dasa (purple; *n*=islets from 3 mice; \*\*\**P*<0.001). (G) Representative normalized (F/F_min_) C57 islet Fura-2 Ca^2+^ responses to 200nM SST and 5μM NSC 87877 in the presence of 20G+1mM Tolb. Insets indicate the percentage of C57 islets exhibiting each type of NSC 87877 [Ca^2+^]_i_ response. (H) Average percentage of C57 islets with 20G+Tolb displaying [Ca^2+^]_i_ oscillations in response to SST (green) and SST+NSC 87877 (pink; *n*=islets from 3 mice; \*\**P*<0.01). (I) Average C57 islet [Ca^2+^]_i_ plateau fraction with 20G+Tolb after SST (green) and after SST+NSC 87877 (pink; *n*=islets from 3 mice; \*\**P*<0.01). (J) Representative normalized (F/F_min_) C57 islet Fura-2 Ca^2+^ responses to 1μM insulin and 100nM Dasa in the presence of 20G+1mM Tolb. (K) Average percentage of C57 islets with 20G+Tolb displaying [Ca^2+^]_i_ oscillations in response to insulin (red) and insulin+Dasa (purple; *n*=islets from 3 mice). (L) Average C57 islet [Ca^2+^]_i_ plateau fraction with 20G+Tolb after insulin (red) and after insulin+Dasa (purple; *n*=islets from 3 mice; \*\**P*<0.01). (M) Immunoblots (IBs) of cell lysates isolated from nontreated C57 islets as well as from islets treated with 200nM SST or 1μM insulin for 15 minutes. IBs were probed for total NKA (ATP1A1), NKA phosphorylated at Y10 (p-ATP1A1^Y10^), and NKA phosphorylated at Y260 (p-ATP1A1^Y260^). Bands corresponding to p-ATP1A1^Y10^ and p-ATP1A1^Y260^ were normalized to total ATP1A1 bands; p-ATP1A1^Y10^ and p-ATP1A1^Y260^ bands from islets with SST and insulin were then normalized to nontreated islets (shown below IBs). Statistical analysis was conducted using unpaired two-sample t-tests (E, F, H, I, and K) or oneway ANOVA (B, C, and L); uncertainty is expressed as mean ± SEM.

It has been shown that SSTR signaling stimulates Src activity (43), which interacts with and enhances NKA function (44, 45, 47). Therefore, a Src inhibitor dasatinib was employed to investigate the role of Src signaling in G_i/o_-coupled GPCR control of β-cell NKA function. All islets displayed SST-induced [Ca^2+^]_i_ oscillations, which decreased to 37.6±9.8% of islets in the presence of 100nM dasatinib (Fig. 6D and 6E; *P*<0.01). Dasatinib also increased islet [Ca^2+^]_i_ plateau fraction by 40.7±3.1% compared to SST alone (Fig. 6D and 6F; *P*<0.001). Interestingly, the average area of islets that oscillated in the presence of dasatinib was 71.0±8.6% larger than islets that stopped oscillating (*P*<0.01). This may indicate that bifurcated islet Ca^2+^ responses are due in-part to poor penetration of dasatinib into larger islets. As SSTR signaling activates Src homology region 2 domain-containing phosphatase-2 (Shp2) that in turn stimulates Src function, a Shp2 inhibitor NSC 87877 was utilized to assess whether this pathway regulates β-cell NKAs. Treatment with 5μM NSC 87877 decreased the percentage of islets displaying SST-induced [Ca^2+^]_i_ oscillations from 100% to 53.5±6.1% (Fig. 6G and 6H; *P*<0.01); islet [Ca^2+^]_i_ plateau fraction also increased by 39.9±6.8% compared to SST alone (Fig. 6G and 6I; *P*<0.01). As with dasatinib treatment, the average area of islets that continued oscillating with NSC 87877 was 75.7±16.7% larger than islets that stopped oscillating (*P*<0.05), which again suggests only partial Shp2 inhibition in larger islets.

Insulin receptor signaling also stimulates Src signaling as well as NKA activity (35, 53). Therefore, we examined whether insulin regulates β-cell NKA function through Src activation. In the presence of 20mM glucose and 1mM tolbutamide exogenous insulin (1μM) induced [Ca^2+^]_i_ oscillations in 69.5±10.4% of islets (Fig. 6J and 6K) and decreased islet [Ca^2+^]_i_ plateau fraction by 16.4±2.6% (Fig. 6J and 6L; *P*<0.01), which was comparable to the effect of SST. Subsequent addition of dasatinib had no effect on the percentage of islets displaying insulin-induced [Ca^2+^]_i_ oscillations (58.6±18.6%; Fig. 6J and 6K). Dasatinib treatment did increase islet [Ca^2+^]_i_ plateau fraction by 6.9±4.0% compared to insulin alone; however, the change was not significant (Fig. 6J and 6L). These results suggest that β-cell NKA function is augmented by insulin receptor signaling, but that this effect is likely not mediated by Src. Furthermore, these findings again suggest that once initiated β-cell NKA function is oscillatory in nature, independent of the mechanism of activation.

Phosphorylation of the NKA α1 subunit by tyrosine kinases has been shown to regulate pump activity (46, 54, 55). Thus, to elucidate the mechanism underlying tyrosine kinase control of β-cell NKA function, mouse islets were treated with SST, insulin, or vehicle; cell lysates were then immunoblotted for total NKA α1 protein as well as for NKA α1 phosphorylated at two putative tyrosine kinase phosphorylation sites (tyrosine 10 (Y10) and tyrosine 260 (Y260)) (46, 54, 56). Phosphorylation of NKA α1 at residue Y10 (p-ATP1A1^Y10^) increased following treatment with SST (1.8-fold increase) and insulin (1.7-fold increase), whereas phosphorylation of NKA α1 at residue Y260 (p-ATP1A1^Y260^) was unaffected (Fig. 6M). As previous studies have shown that phosphorylation of NKA α1 at residue Y10 increases NKA activity (55), this result suggests that tyrosine kinases augment β-cell NKA function in-part through direct phosphorylation of the NKA α1 subunit.

## DISCUSSION

Islet G_i/o_-coupled GPCR signaling modulates β-cell [Ca^2+^]_i_ oscillations, which in turn regulates pulsatile insulin secretion (6, 18, 21). Here, we elucidate the mechanistic underpinnings of β-cell G_i/o_ signaling-induced *V*_m_ hyperpolarization. We found that G_i/o_ signaling activates β-cell NKAs, which controls the frequency of islet [Ca^2+^]_i_ oscillations and initiates [Ca^2+^]_i_ oscillations independently of K_ATP_ channel activity. Furthermore, Src activation was required for G_i/o_ signaling-induced stimulation of β-cell NKAs; however, NKA-mediated *V*_m_ hyperpolarization was also inhibited by cAMP-dependent PKA activation, indicating that multiple signaling modalities converge to control NKA function and thus islet [Ca^2+^]_i_ handling. Therefore, these results demonstrate the importance of δ-cell SST secretion in controlling islet [Ca^2+^]_i_ oscillations through SSTR-mediated regulation of β-cell NKA activity. Moreover, we have identified a conserved mechanism for G_i/o_-coupled GPCR control of β-cell NKA function that extends to ADRs and DRDs. Therefore, the data reveal that G_i/o_-coupled GPCR control of β-cell NKA activity plays a critical role in tuning the frequency of islet [Ca^2+^]_i_ oscillations, and thus the kinetics of insulin secretion.

Activation of GIRK channels accounts for G_i/o_ signaling-induced *V*_m_ hyperpolarization In many tissues (37); this has also been proposed for β-cells (16, 36). However, biophysical evidence of β-cell GIRK channel activation though G_i/o_-coupled GPCR signaling is limited. Indeed, elegant studies by Sieg et al. demonstrated that GIRK channels are not activated by ADR stimulation (19). We also found no evidence of an SSTR-mediated GIRK conductance in mouse or human β-cells. Moreover, SST-activated currents were outward at voltages below the equilibrium potential of K^+^, were present in β-cells without GIRK2 channels, and were not inhibited by tertiapin-Q. Nevertheless, as GIRK channels are expressed in β-cells and SST-induced currents trended lower in the presence of tertiapin-Q, a role for these channels cannot be completely discounted. However, the primary mechanism responsible for G_i/o_ signaling-induced inhibition of insulin secretion has remained elusive for more than 50 years. As many studies have shown that G_i/o_ signaling hyperpolarizes β-cell *V*_m_ (16, 18, 19, 36), it was assumed that a G_i/o_-coupled GPCRs activate a K^+^ channel. To enhance K^+^ currents through this putative channel, K^+^ was removed from the extracellular solution, but surprisingly, SST-induced β-cell currents and *V*_m_ hyperpolarization were both abolished. Interestingly, in the absence of extracellular K^+^, diazoxide was still able to mediate a decrease in islet [Ca^2+^]_i_, which suggested that SST-induced *V*_m_ hyperpolarization was not due to a K^+^ current. As G_i/o_-coupled GPCR signaling activates NKAs in other tissues and removal of extracellular K^+^ inhibits NKA activity, we considered NKAs a strong candidate to explain SST-induced β-cell *V*_m_ hyperpolarization (38, 40). This was confirmed using the NKA inhibitor ouabain, which blocked SST-induced reductions of islet [Ca^2+^]_i_. Furthermore, epinephrine and dopamine have been shown to hyperpolarize β-cell *V*_m_ and stimulate NKAs in certain tissues (8–10, 16, 18, 38), and indeed, our data confirmed that α2A-adrenergic signaling regulates β-cell [Ca^2+^]_i_ by activating NKAs. Taken together, these results suggest that activation of NKAs rather than K^+^ channels serves as a conserved mechanism for G_i/o_ signaling-mediated β-cell *V*_m_ hyperpolarization.

The mechanisms underlying G_i/o_-coupled GPCR control of NKA activity are highly tissue-specific, but are predominantly linked to the phosphorylation status of the NKA α subunit (38, 39, 46). Our data support previous findings showing that G_i/o_ signaling-induced islet [Ca^2+^]_i_ oscillations can be terminated by stimulating adenylyl cyclase activity, inhibiting phosphodiesterase activity, or directly raising [cAMP]_i_ (21); this suggests that phosphorylation by cAMP-dependent kinases inhibits β-cell NKAs. Furthermore, as both glucose metabolism and Gs-coupled GPCR signaling increase β-cell [cAMP]_i_ levels (52, 57–60), this signal would be predicted to dynamically regulate NKA activity and thus Ca^2+^ handling. Indeed, several minutes after SST treatment we observed robust [cAMP]_i_ oscillations that were out of phase with [Ca^2+^] oscillations. Pharmacological blockade of PKA signaling also decreased islet [Ca^2+^]_i_ and prevented forskolin-mediated inhibition of β-cell NKAs. Therefore, these findings suggest that G_i/o_ signaling-induced fluctuations in islet [cAMP]_i_ influence oscillations in β-cell NKA activity. While PKA is the primary cAMP-regulated β-cell kinase, additional kinases that modulate NKA function (i.e. PKC, EPAC, PKG) could also be impacted by changes in [cAMP]_i_ (61–65). Thus, future studies are required to further determine how [cAMP]_i_ tunes NKA activity not only via G_i/o_-coupled GPCR signaling, but also through pathways that increase [cAMP]_i_ such as glucose metabolism and Gs-coupled GPCRs.

Although islet [cAMP]_i_ oscillations were observed subsequent to [Ca^2+^] oscillations, SST treatment caused only a very modest initial reduction in islet [cAMP]_i_; this suggests that cAMP is not the principal signal controlling G_i/o_-coupled GPCR-induced islet [Ca^2+^] oscillations. While cAMP-dependent changes in PKA activity modulate β-cell NKA function, G_i/o_-coupled GPCRs also stimulate cAMP-independent protein kinases that have been shown to influence NKA activity. For example, the tyrosine kinase Src and the tyrosine phosphatase Shp2, which activates Src, interact with SSTRs and are activated by SSTR signaling (43, 66). Moreover, Src also interacts with and augments NKA activity (44, 47). As Src and Shp2 inhibition greatly diminished NKA-mediated decreases of islet [Ca^2+^]_i_, SST-induced Src signaling is predicted to facilitate activation of β-cell NKAs. Interestingly, the Src inhibitor used in these studies, dasatinib, which is FDA approved for the treatment of Philadelphia chromosome-positive chronic myeloid leukemia, has been shown to decrease blood glucose levels in numerous clinical studies, most prominently in diabetic patients (67–69). This suggests the possibility that dasatinib increases human islet insulin secretion by inhibiting NKAs, and thus enhancing glucose-mediated Ca^2+^ entry. Insulin receptors that interact with and control Src signaling are also tyrosine kinases and have been shown to stimulate NKAs in a variety of tissues (35, 53, 70). Our findings confirmed that insulin enhances β-cell NKA function; however, insulin-induced islet [Ca^2+^]_i_ oscillations were only modestly affected by dasatinib treatment. This indicates that insulin stimulates β-cell NKAs independently of Src signaling, presumably through direct phosphorylation of NKA α1 subunits. Both insulin and Src have been shown to phosphorylate NKA α1 at residue Y10, which augments NKA function (46, 55, 56). This is supported by the finding that phosphorylation of NKA α1 residue Y10 increases in islets following treatment with SST or insulin, which indicates that β-cell NKA activation is in-part due to phosphorylation of the NKA α1 subunit by tyrosine kinases. Therefore, it will be important to further examine the mechanisms underlying tyrosine kinase regulation of β-cell NKA function and its role in tuning islet [Ca^2+^]_i_ oscillations as well as GSIS.

SST secretion becomes defective during the pathogenesis of diabetes (33, 71); therefore, control of β-cell NKA function by SSTR signaling would also be expected to be perturbed during T2D. Other islet cell types, including α- and δ-cells, also express high levels of NKA α1 subunit transcript (11–13), which is supported by our immunofluorescence staining of human pancreatic sections that showed NKA α1 subunit expression in insulin-negative islet cells. Furthermore, numerous NKA β and γ subunit transcripts as well as several G_i/o_-coupled GPCRs are expressed in α-cells (i.e. SST, D2-like DRDs, and α1-ADRs) and δ-cells (i.e. D2-like DRDs and α2-ADRs) (11–13, 16, 30, 72). Moreover, NKAs have been shown to regulate α-cell *V*_m_ in response to fatty acid metabolism (73). Thus, NKA control of plasma membrane *V*_m_ would be expected to influence [Ca^2+^]_i_ dynamics in these other islet cell types and may in-part explain why GIRK channel inhibition fails to completely inhibit SST-induced α-cell *V*_m_ hyperpolarization (49). Lastly, expression of specific NKA subunits, such as FXYD2, are altered in T2D human islets (11, 74) and in leptin receptor deficient diabetic mouse islets (75). Thus, understanding the role(s) that NKA serves in all islet cell types under physiological and diabetic conditions will illuminate critical features of islet function and disfunction.

In summary, we identified a conserved G_i/o_-coupled mechanism for controlling β-cell Ca^2+^ entry, and thus insulin secretion in response to numerous G_i/o_-coupled GPCR ligands that have been shown to limit insulin secretion (i.e. adrenaline, SST, and dopamine). Moreover, we demonstrated that endogenous SSTR signaling tunes islet [Ca^2+^]_i_ oscillations and presumably pulsatile insulin secretion by activating β-cell NKAs. Finally, we determined that G_i/o_ signaling stimulates β-cell NKA function by activating tyrosine kinases (e.g. Src and insulin receptors). Taken together, these findings reveal an essential and conserved NKA-mediated mechanism governing G_i/o_-coupled signals known to regulate insulin secretion that remained elusive for over half a century (Fig. 7).

**Fig. 7:**
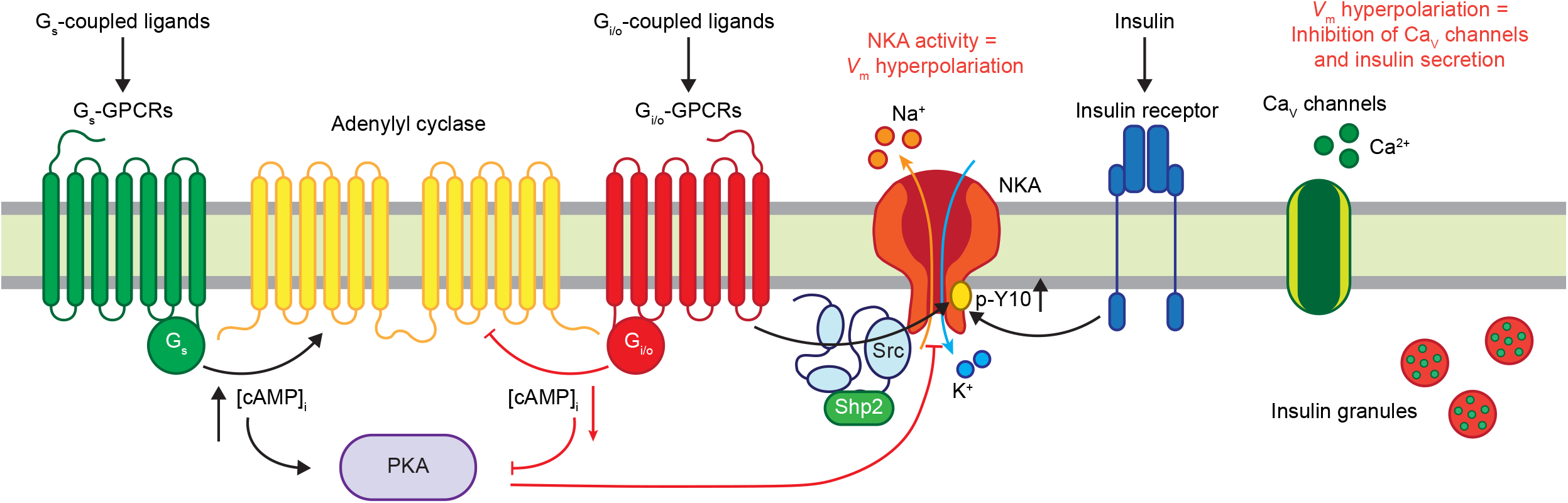
Model illustrating the mechanisms that regulate β-cell NKA function. (A) Overview of stimulatory and inhibitory receptor-mediated signaling pathways that tune β-cell NKA activity. G_i/o_-coupled GPCR signaling transiently hyperpolarizes β-cell *V*_m_ via Src-mediated phosphorylation of NKAs as well as by decreasing [cAMP]_i_ and PKA activity. Other tyrosine kinases (e.g. insulin receptors) also phosphorylate and activate β-cell NKAs. Stimulation of Gs-coupled GPCRs is predicted to limit β-cell NKA function by increasing [cAMP]_i_ and PKA activity. This figure was created with BioRender.com.

## METHODS

### Animals

All mice were 12-to 18-week old, age-matched males on a C57Bl6/J background (Stock #: 000664; The Jackson Laboratory (JAX), Bar Harbor, ME). Transgenic mice expressing G_i/o_-coupled protein-coupled Designer Receptors Exclusively Activated by Designer Drugs (DREADDs) specifically in δ-cells (δGi DREADDs) were generated by crossing mice expressing a mutant G_i/o_-coupled GPCR P2A mCitrine fluorescent reporter (B6.129-Gt(ROSA)^26Sortm1(CAG-CHRM4*,-mCitrine)Ute/J^; Stock #: 026219; JAX) construct preceded by a loxP-flanked STOP cassette with mice expressing a *SST*-IRES-Cre (Stock #: 013044; JAX) (76, 77). All δGi DREADD islets were prepared from mice heterozygous for *SST*-IRES-Cre as endogenous SST is decreased in an allele dosage-dependent manner. Transgenic mice with pancreatic-specific knockout (KO) of *Kcnj6* (gene encoding GIRK2) were generated by crossing animals with a loxP-flanked *Kcnj6* exon 4 with mice expressing a *Pdx1-Cre* (GIRK2 KO^Panc^; Stock #: 014647; JAX) (78, 79). All animals were housed in a Vanderbilt University IACUC (protocol # M1600063-01) approved facility on a 12-hour light/dark cycle with access to standard chow (Lab Diets, 5L0D) *ad libitum.* Mice were humanely euthanized by cervical dislocation followed by exsanguination. To preserve islet ion channel function mice were not treated with anesthesia.

### Human donors

All studies detailed here were approved by the Vanderbilt University Health Sciences Committee Institutional Review Board (IRB# 110164). Healthy human islets were provided from multiple isolation centers by the Integrated Islet Distribution Program (IIDP). Deidentified human donor information is provided in Supplemental Table 1. The IIDP obtained informed consent for deceased donors in accordance with NIH guidelines prior to reception of human islets for our studies.

### Chemicals and reagents

Unless otherwise noted all chemicals and reagents were purchased from Sigma-Aldrich (St. Louis, MO) or Thermo Fisher (Waltham, MA). Clozapine N-oxide (CNO) was purchased from Hello Bio (Princeton, NJ). Clonidine hydrochloride (Clon), forskolin (FSK), and ouabain (Oua) were purchased from R&D Systems (Minneapolis, MN). Tertiapin-Q (TPQ) was purchased from Alomone Labs (Jerusalem, Israel). Dasatinib (Dasa), H-89, and NSC 87877 were purchased from Cayman Chemical (Ann Arbor, MI). An optimized rat insulin promoter (RIP) (50) and the coding sequence of hM4D(Gi)-mCherry (G_i/o_-coupled DREADD-mCherry fusion protein; Plasmid #75033; Addgene, Watertown, MA) (80) were cloned into a pLenti6 lentiviral transfer plasmid and utilized to produce 3rd-generation lentiviruses (LVs) as previously described (81).

### Islet isolation

Mouse pancreata were digested with collagenase P (Roche; Basel, Switzerland) and islets were isolated using density gradient centrifugation (82); mouse islets were cultured in RPMI-1640 (Corning) media with 5.6 mM glucose supplemented with 15% FBS, 100 IU·ml^-1^ penicillin, and 100mg·ml^-1^ streptomycin (RPMI) at 37°C, 5% CO_2_. Upon arrival, human islets were allowed to recover for at least 2 hours in CMRL-1066 (Corning, Cleveland, TN) media containing 5.6mM glucose and supplemented with 20% fetal bovine serum (FBS), 100IU·ml^-1^ penicillin, 100mg·ml^-1^ streptomycin, 2mM GlutaMAX, 2mM HEPES, and 1mM sodium pyruvate (CMRL) at 37°C, 5% CO_2_. Mouse and human islets were cultured in poly-d-lysine-coated 35mm glass-bottomed dishes (CellVis, Mountain View, CA) and all experiments were conducted within 48 hours.

### Patch-clamp electrophysiology

Patch electrodes (3-4MΩ) were backfilled with intracellular solution containing (mM) 90.0 KCl, 50.0 NaCl, 1.0 MgCl_2_, 10.0 EGTA, 10.0 HEPES, and 0.005 amphotericin B (adjusted to pH 7.2 with KOH). Mouse and human islets were patched in Krebs-Ringer HEPES buffer (2mL; KRHB) containing (mM) 119.0 NaCl, 4.7 KCl, 2.0 CaCl_2_, 1.2 MgSO_4_, 1.2 KH_2_PO_4_, and 10.0 HEPES (pH 7.35 adjusted by NaOH) supplemented with 2mM glucose; a perforated whole-cell patch-clamp technique was utilized to record β-cell membrane potential (*V*_m_) in current-clamp mode using an Axopatch 200B amplifier with pCLAMP10 software (Molecular Devices). Islet cells that did not display electrical activity at 2mM glucose were identified as β-cells. After a perforated patch configuration was established (seal resistance>1.0GΩ; leak<20.0pA) *V*_m_ depolarization and action potential (AP) firing were induced by exchanging the bath solution with KRHB supplemented with 20mM glucose and 1mM tolbutamide. After AP firing was observed, the amplifier was switched to voltage-clamp mode; *V*_m_ was held at −60mV and ramped from −100mV to −50mV every 15 seconds for at least 2 minutes and the resulting β-cell currents were recorded. The amplifier was then returned to current-clamp mode and *V*_m_ was recorded. Mouse and human islets were perifused with further treatments and changes in β-cell *V*_m_ and currents were measured as indicated in figure legends.

### Intracellular Ca^2+^ and cAMP imaging

For simultaneous islet [Ca^2+^]_i_ and [cAMP]_i_ imaging, mouse islets were transduced for 4 hours with an adenovirus (AV) expressing a CMV-jRGECO1a-P2A-cAMPr construct (VectorBuilder, Chicago, IL) and cultured 24 hours at 37°C, 5% CO_2_ prior to imaging. Alternatively, mouse islets were loaded with a Fura-2 AM Ca^2+^ indicator (2μM) for 30 minutes before the start of an experiment. Human islets were either transduced for 4 hours with an AV expressing RIP-GCaMP6s (VectorBuilder) and cultured 48 hours before imaging or loaded with a Cal-590 Ca^2+^ indicator (10μM; AAT Bioquest, Sunnyvale, CA) for 1 hour prior to the start of a study. Before each experiment, islet culture media was replaced with 2mL KRHB supplemented with 2mM glucose; after 10 minutes, the islets were treated as detailed in figure legends. Mouse islet jRGECO1a (excitation (Ex): 561nm; emission (Em): 620±50nm) and cAMPr fluorescence (Ex: 488nm; Em: 525±32nm) were simultaneously measured every 5 seconds utilizing an LSM 780 multi-photon confocal microscope equipped with Zeiss Zen software (20x magnification; LSM 780) as indicators of [Ca^2+^]_i_ and [cAMP]_i_ respectively. Mouse islet Fura-2 AM fluorescence (Ex: 340nm and 380nm; Em: 510±40nm) was measured every 5 seconds with a Nikon Ti2 epifluorescence microscope equipped with a Prime 95B camera with 25mm CMOS sensors and Nikon Elements software (10x magnification; Nikon Ti2); the ratio of Fura-2 AM fluorescence excited at 340nm and 380nm was utilized as an indicator of [Ca^2+^]_i_. Human islet Cal-590 (Ex: 560±20nm; Em: 630±37.5nm) or β-cell GCaMP6s fluorescence (Ex: 488nm; Em: 531±48nm) was measured every 5 seconds as an indicator of [Ca^2+^]_i_ utilizing the LSM 780 microscope (20x magnification).

### Intracellular Ca^2+^ and Na^+^ imaging

For simultaneous islet [Ca^2+^]_i_ and [Na]i imaging, mouse islets were loaded with a Fura Red AM Ca^2+^ indicator (5μM; catalog #: F3021; Thermo Fisher) and an ION Natrium Green-2 (ING-2) Na^+^ indicator (5μM; catalog #: 2011F; Ion Biosciences, San Marcos, TX) for 1 hour before the start of an experiment. Before each experiment, islet culture media was replaced with KRHB supplemented with 2mM glucose; after 10 minutes, the islets were treated as detailed in figure legends. Mouse islet Fura Red (Ex: 430±12nm and 500±10nm; Em: 700±37.5nm) fluorescence was measured every 5 seconds with the Nikon Ti2 (10x magnification); the ratio of Fura Red fluorescence excited at 500±10nm and 430±12nm was utilized as an indicator of [Ca^2+^]_i_. Mouse islet ING-2 fluorescence (Ex: 500±10nm; Em: 535±15nm) was measured simultaneously every 5 seconds as an indicator of [Na]_i_.

### Immunofluorescence imaging

Paraffin-embedded human pancreas sections were processed and probed as previously described (81). Following rehydration, sections were subject to Tris-EDTA-SDS antigen retrieval at 37°C for 40 minutes (83); pancreas sections were stained with primary antibodies (1:100 mouse anti-ATP1A1 (catalog #: MA3-928; Thermo Fisher) and 1:1000 guinea pig anti-insulin (catalog #: 20-IP35; Fitzgerald, North Acton, MA) followed by secondary antibodies (1:300 donkey antimouse Alexa Fluor 647 (catalog #: 715-606-150; Jackson ImmunoResearch, West Grove, PA) and 1:300 donkey anti-guinea pig Alexa Fluor 488 (catalog #: 706-546-148; Jackson ImmunoResearch)). Immunofluorescence images were collected with a Zeiss LSM 710 META inverted confocal microscope (40x magnification).

### Hormone secretion assays

Mouse islets were cultured overnight in RPMI (supplemented with 0.5mg/mL BSA) then transferred to equilibration media (DMEM (no glucose) with 10% FBS, 0.5mg/mL BSA, 10mM HEPES, and 0.5mM CaCl_2_) supplemented with 5.6mM glucose for 1 hour at 37°C, 5% CO_2_. Islets were picked on ice into a 24-well plate (Corning) containing 400μL of secretion media (DMEM (no glucose) with 0.5mg/mL BSA, 10mM HEPES, and 0.5mM CaCl_2_) supplemented with the glucose concentrations and treatments indicated in figure legends then cultured for 1 hour at 37°C, 5% CO_2_. Secretion was halted by transferring the plate to ice for 10 minutes and supernatants were collected in low retention 1.6mL centrifuge tubes. Supernatants were supplemented with 1:100 mammalian protease inhibitor cocktail and stored at −20°C until analyzed. Secreted insulin was measured as per manufacturer instructions with ALPCO insulin chemiluminescence ELISA kits (15 islets per sample; catalog #: 80-INSHU-CH01) or Mercodia mouse insulin ELISA kits (50 islets per sample; catalog #: 10-1247-01); secreted SST was measured as per manufacturer instructions with Phoenix Pharmaceuticals SST chemiluminescent EIA kits (50 islets per sample; CEK-060-03).

### NKA immunoblotting

Mouse islets were isolated and incubated overnight in RPMI supplemented with 5.6mM glucose at 37°C and 5% CO_2_. The islets were transferred to KRHB supplemented with treatments indicated in figure legends for 15 minutes at room temperature, washed with phosphate-buffered saline supplemented with identical treatments along with 20μL/mL Halt protease/phosphatase inhibitor cocktail (Halt PPI; catalog #: 78442), and frozen in a dry ice ethanol bath. Islets were lysed on ice in RIPA buffer supplemented with Halt PPI, then cell lysates were resolved on nitrocellulose membranes. Immunoblots were blocked for 1 hour in Tris-buffered saline with 0.1% Tween 20 (TBST) supplemented with 5% BSA. All primary and secondary antibodies were diluted in TBST supplemented with 0.1 % BSA. Immunoblots were probed with 1:500 rabbit anti-phospho-ATP1A1 (Y10) (p-ATP1A1^Y10^; catalog #: PA5-17061; Thermo Fisher) followed by 1:2500 goat anti-rabbit HRP-conjugated secondary (catalog #: W4011; Promega, Madison, WI). p-ATP1A1^Y10^ protein bands were visualized with SuperSignal™ West Pico Plus (SuperSignal Pico; catalog #: 34580; Thermo Fisher) utilizing a Bio-Rad Digital ChemiDoc MP (ChemiDoc). Immunoblots were stripped for 20 minutes with Restore™ Western Blot Stripping Buffer (catalog #: 21059; Thermo Fisher) and re-probed with the following: 1:500 rabbit anti-phospho-ATP1A1 (Y260) (p-ATP1A1^Y260^; catalog #: PA5-118769; Thermo Fisher) followed by 1:2500 goat anti-rabbit HRP-conjugated secondary; 1:1000 mouse anti-ATP1A1 followed by 1:2500 goat anti-mouse HRP-conjugated secondary (catalog #: W4021; Promega). p-ATP1A1^Y260^ and ATP1A1 bands were visualized with SuperSignal Pico utilizing the ChemiDoc.

### Data analysis

Islet [Ca^2+^]_i_, [cAMP]_i_, [Na^+^]_i_, and NKA immunofluorescence were analyzed using Zeiss Zen software, Nikon Elements software, and the ImageJ Fiji image processing pack. Axon Clampfit software was utilized to quantify β-cell SST-induced currents and *V*_m_ as well as perform crosscorrelation analyses. All β-cell SST-induced currents are median values of 10 or more consecutive traces. Heatmaps were generated using the MATLAB imagesc function. Period analysis of δGi islet [Ca^2+^]_i_ oscillations was carried out utilizing the MATLAB detrend and findpeaks functions. Immunoblots were analyzed using Bio-Rad Image Lab 5.0. Figures were prepared utilizing Adobe Illustrator. Statistical analyses were carried out utilizing Microsoft Excel and GraphPad Prism 9.2.0 as indicated in figure legends; data were compared utilizing paired or unpaired two-sample t-tests, one-sample t-tests, or one-way analysis of variance (ANOVA) with Šidák’s post-hoc multiple comparisons tests. Data were normalized when appropriate as indicated in figure legends. Unless stated otherwise, data are presented as mean values ± standard error (SEM) for the specified number of samples (*n*). Differences were considered significant for *P*≤0.05.

### Study Approval

Animal: All animals employed in these studies were handled in compliance with guidelines approved by the Vanderbilt University Institutional Animal Care and Use Committee (IACUC). Human: All studies detailed here were approved by the Vanderbilt University Health Sciences Committee Institutional Review Board (IRB# 110164). The IIDP obtained informed consent for deceased donors in accordance with NIH guidelines prior to reception of human islets for these studies.

## AUTHOR CONTRIBUTIONS

MTD and DAJ formulated and designed experiments. MTD, PKD, KEZ, AYN, CMS, JRD, NMW, JCL, and CS performed experiments. MTD and DAJ analyzed data. MTD and DAJ interpreted experimental results. MTD and DAJ prepared figures. MTD and DAJ drafted the manuscript. MTD, PKD, KEZ, AYN, CMS, JRD, NMW, JCL, CS, and DAJ approved the final manuscript submitted for publication.

## ACKNOWLEDGEMENTS

We thank Dr. Kevin D. Wickman (University of Minnesota) for providing mice with a loxP-flanked Kcnj6 exon 4. We are also grateful for the confocal microscopy resources provided by the Vanderbilt Cell Imaging Shared Resource (CISR; supported by NIH grants CA68485, DK20593, DK58404, DK59637 and EY08126). This research was performed with the support of the Integrated Islet Distribution Program (https://iidp.coh.org/). We especially thank the organ donors and their families.

---

**Fig. S1:**
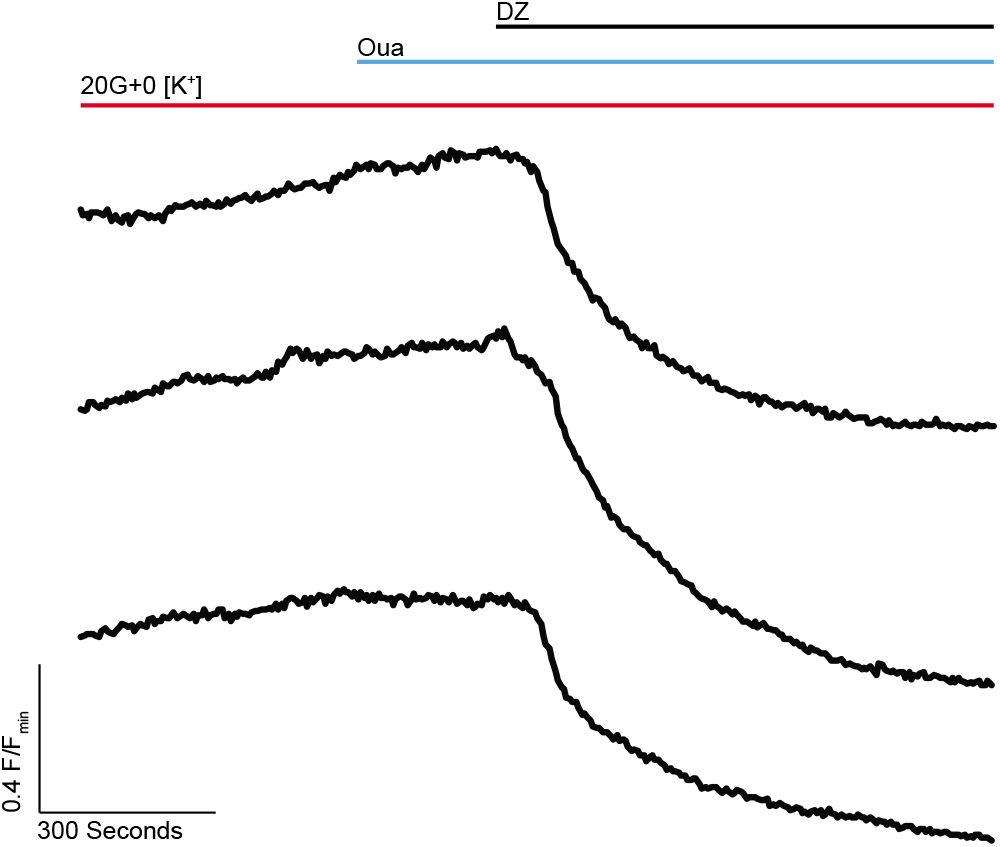
K_ATP_ channel activation can hyperpolarize β-cell *V*_m_ when NKAs are inactive. Representative normalized (F/F_min_) Fura-2 Ca^2+^ responses to 125μM DZ in the presence of 0 [K^+^] and 125μM Oua (*n*=islets from 3 mice). Data analysis is presented in Fig. 2E.

